# Large-Scale Uniform Analysis of Cancer Whole Genomes in Multiple Computing Environments

**DOI:** 10.1101/161638

**Authors:** Christina K. Yung, Brian D. O’Connor, Sergei Yakneen, Junjun Zhang, Kyle Ellrott, Kortine Kleinheinz, Naoki Miyoshi, Keiran M. Raine, Romina Royo, Gordon B. Saksena, Matthias Schlesner, Solomon I. Shorser, Miguel Vazquez, Joachim Weischenfeldt, Denis Yuen, Adam P. Butler, Brandi N. Davis-Dusenbery, Roland Eils, Vincent Ferretti, Robert L. Grossman, Olivier Harismendy, Youngwook Kim, Hidewaki Nakagawa, Steven J. Newhouse, David Torrents, Lincoln D. Stein, on behalf of the PCAWG Technical Working Group, Javier Bartolomé Rodriguez, Keith A. Boroevich, Rich Boyce, Angela N. Brooks, Alex Buchanan, Ivo Buchhalter, Niall J. Byrne, Andy Cafferkey, Peter J. Campbell, Zhaohong Chen, Sunghoon Cho, Wan Choi, Peter Clapham, Francisco M. De La Vega, Jonas Demeulemeester, Michelle T. Dow, Lewis J. Dursi, Juergen Eils, Claudiu Farcas, Francesco Favero, Nodirjon Fayzullaev, Paul Flicek, Nuno A. Fonseca, Josep L.l. Gelpi, Gad Getz, Bob Gibson, Michael C. Heinold, Julian M. Hess, Oliver Hofmann, Jongwhi H. Hong, Thomas J. Hudson, Daniel Huebschmann, Barbara Hutter, Carolyn M. Hutter, Seiya Imoto, Sinisa Ivkovic, Seung-Hyup Jeon, Wei Jiao, Jongsun Jung, Rolf Kabbe, Andre Kahles, Jules Kerssemakers, Hyunghwan Kim, Hyung-Lae Kim, Jihoon Kim, Jan O. Korbel, Michael Koscher, Antonios Koures, Milena Kovacevic, Chris Lawerenz, Ignaty Leshchiner, Dimitri G. Livitz, George L. Mihaiescu, Sanja Mijalkovic, Ana Mijalkovic Lazic, Satoru Miyano, Hardeep K. Nahal, Mia Nastic, Jonathan Nicholson, David Ocana, Kazuhiro Ohi, Lucila Ohno-Machado, Larsson Omberg, B.F. Francis Ouellette, Nagarajan Paramasivam, Marc D. Perry, Todd D. Pihl, Manuel Prinz, Montserrat Puiggròs, Petar Radovic, Esther Rheinbay, Mara W. Rosenberg, Charles Short, Heidi J. Sofia, Jonathan Spring, Adam J. Struck, Grace Tiao, Nebojsa Tijanic, Peter Van Loo, David Vicente, Jeremiah A. Wala, Zhining Wang, Johannes Werner, Ashley Williams, Youngchoon Woo, Adam J. Wright, Qian Xiang, the PCAWG Network

## Abstract

The International Cancer Genome Consortium (ICGC)’s Pan-Cancer Analysis of Whole Genomes (PCAWG) project aimed to categorize somatic and germline variations in both coding and non-coding regions in over 2,800 cancer patients. To provide this dataset to the research working groups for downstream analysis, the PCAWG Technical Working Group marshalled ~800TB of sequencing data from distributed geographical locations; developed portable software for uniform alignment, variant calling, artifact filtering and variant merging; performed the analysis in a geographically and technologically disparate collection of compute environments; and disseminated high-quality validated consensus variants to the working groups. The PCAWG dataset has been mirrored to multiple repositories and can be located using the ICGC Data Portal. The PCAWG workflows are also available as Docker images through Dockstore enabling researchers to replicate our analysis on their own data.

## Introduction

The International Cancer Genome Consortium (ICGC)/The Cancer Genome Atlas (TCGA) Pan-Cancer Analysis of Whole Genomes (PCAWG) study has characterized the pattern of mutations in over 2,800 cancer whole genomes. Extending TCGA Pan-Cancer analysis project, which focused on molecular aberrations in protein coding regions only^1^, PCAWG undertook the study of whole genomes, allowing for the discovery of driver mutations in cis-regulatory sites and non-coding RNAs, examination of the patterns of large-scale structural rearrangements, identification of signatures of exposure, and elucidation of interactions between somatic mutations and germline polymorphisms.

The PCAWG dataset comprises a total of 5,789 whole genomes of tumors and matched normal tissue spanning 39 tumor types. The tumor/normal pairs came from a total of 2,834 donors collected and sequenced by 48 sequencing projects across 14 jurisdictions (Supplementary Fig. 1). In addition, RNA-Seq profiles were obtained from a subset of 1,284 of the donors^2^. While the individual sequencing projects contributing to PCAWG had previously identified genomic variants within their individual cancer cohorts, each project had used their own preferred methods for read alignment, variant calling and artifact filtering. During initial evaluation of the data set, we found that the different analysis pipelines contributed high levels of technical variation, hindering comparisons across multiple cancer types^3^. To eliminate the variations arising from non-uniform analysis, we reanalyzed all samples starting with the raw sequencing reads and using a standardized set of alignment, variant calling and filtering methods. These “core” workflows yielded uniformly analyzed genomic variants for downstream analyses by various PCAWG working groups. A subset of these variants were validated through targeted deep sequencing to estimate the accuracy of our approach^4^.

To create this uniform analysis set, multiple logistic and technical challenges had to be overcome. First, projects participating in the PCAWG study employed their own metadata conventions for describing their raw sequencing data sets. Hence, we had to establish a PCAWG metadata standard suitable for all the participating projects. Second, and more significantly, the data was large in size -- 800TB of raw sequencing reads -- and distributed geographically across the world. During realignment, the data transiently doubled in size, and after final variant calling and other downstream analysis, the full data set reached nearly 1PB. Furthermore, the compute necessary to fully harmonize the data was estimated at more than 30 million core-hours. Both the storage and compute requirements made it impractical to complete the analysis at any single research institute. In addition, legal constraints across the various jurisdictions imposed restrictions as to where personal data could be stored, analyzed and redistributed^5^. Hence, we needed a protocol to spread the compute and storage resources across multiple commercial and academic compute centers. This requirement, in turn, necessitated the development of analysis pipelines that would be portable to different compute environments and yield consistent analysis results independent of platform. With multiple analysis pipelines running simultaneously in multiple compute environments, the assignment of workload, tracking of progress, quality checking of data and dissemination of results all required sophisticated and flexible planning.

Our approach to tackling these challenges was unique and substantially different from previous large-scale genome analysis endeavors. First, as a collaborative effort among a wide range of institutions not backed by a centralized funding source, a high degree of coordination among a large task force of volunteer software engineers, bioinformaticians and computer scientists was required. Second, the project fully embraced the use of both public and private cloud compute technologies while leveraging established high-performance computing (HPC) infrastructures to fully utilize the compute resources contributed by the partner organizations. The cloud technology platforms we utilized included both Infrastructure as a Service (IaaS): OpenStack, Amazon Web Services and Microsoft Azure; and Platform as a Service (PaaS): Seven Bridges (SB). Lastly, the project made heavy use of Docker, a new lightweight virtualization technology that ensured workflows, tools and infrastructure would work identically across the large number of compute environments utilized by the project.

Utilizing the compute capacity contributed by academic HPC, academic clouds and commercial clouds (Table 1), we were able to complete a uniform analysis of the entire set of 5,789 whole genomes in just over 23 months (Figure 1). Figure 3 illustrates the three broad phases of the project: (1) Marshalling and upload of the data into data analysis centres (3 months); (2) Alignment and variant calling (18 months); and (3) Quality filtering, merging, synchronization and distribution of the variant calls to downstream research groups (2 months). A fourth phase of the project, in which PCAWG working groups used the uniform variant calls for downstream analysis, such as cancer driver discovery, began in the summer of 2016 and continued through the first two quarters of 2017.

**Table 1.**
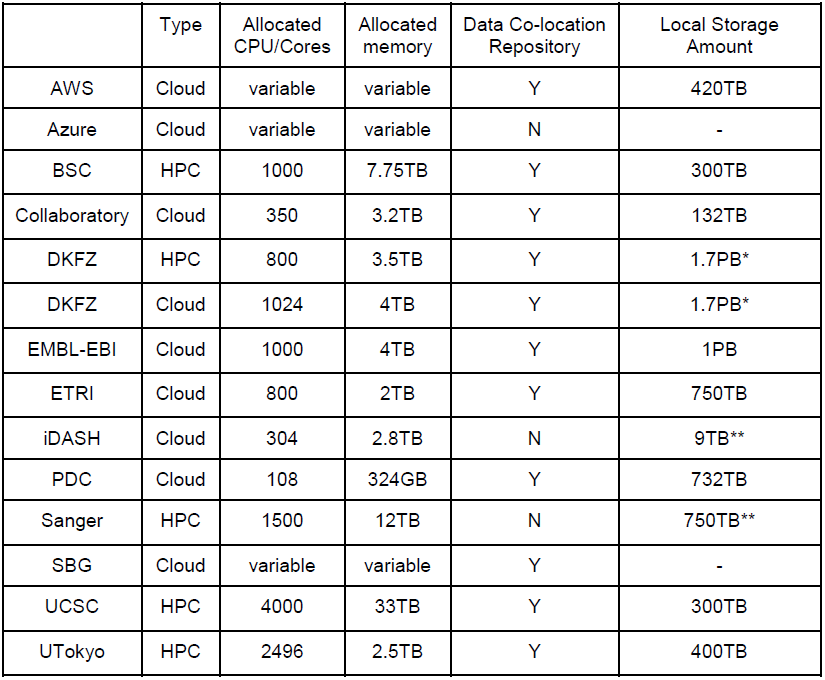
Compute resources. * Shared between environments. ** Transient storage used for local data processing.

|  | Type | Allocated CPU/Cores | Allocated memory | Data Co-location Repository | Local Storage Amount |
| --- | --- | --- | --- | --- | --- |
| AWS | Cloud | variable | variable | Y | 420TB |
| Azure | Cloud | variable | variable | N | - |
| BSC | HPC | 1000 | 7.75TB | Y | 300TB |
| Collaboratory | Cloud | 350 | 3.2TB | Y | 132TB |
| DKFZ | HPC | 800 | 3.5TB | Y | 1.7PB* |
| DKFZ | Cloud | 1024 | 4TB | Y | 1.7PB* |
| EMBL-EBI | Cloud | 1000 | 4TB | Y | 1PB |
| ETRI | Cloud | 800 | 2TB | Y | 750TB |
| iDASH | Cloud | 304 | 2.8TB | N | 9TB** |
| PDC | Cloud | 108 | 324GB | Y | 732TB |
| Sanger | HPC | 1500 | 12TB | N | 750TB** |
| SBG | Cloud | variable | variable | Y | - |
| UCSC | HPC | 4000 | 33TB | Y | 300TB |
| UTokyo | HPC | 2496 | 2.5TB | Y | 400TB |

**Figure 1:**
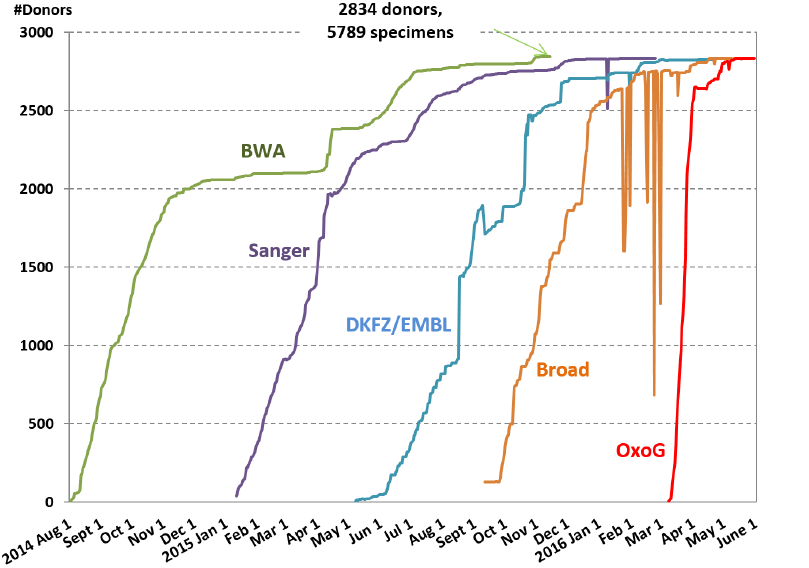
Progress of the 5 workflows over time. The “flat line” of the BWA workflow was due to two major tranches of sequencing data submissions, with a first tranche of ~2000 donors and a second tranche of ~800 donors that were uploaded later. The staggered start of the three variant calling pipelines was dictated more by the time required to develop and package the workflows, and less by the availability of compute power. The “dips” on the plots resulted from quality issues with some sets of variant calls that were withdrawn, reprocessed and resubmitted. In the case of the Broad workflow, the variant calls were withdrawn for post-processing before being considered complete. If all workflows and data would have been in place at the beginning of the project, we estimate the computation across the full set of 5,789 genomes could have been completed in under 6 months.

The following sections will describe the technical solutions used to accomplish each of the phases of the project.

## Phase 1: Data Marshalling and Upload

A significant challenge for the project was that at its inception, a large portion of the raw read sequencing data had yet to be submitted to a read archive and thus had no standard retrieval mechanism. In addition, the metadata standards for describing the raw data varied considerably from project to project. For this reason, we asked the participating projects to prepare and upload the 774 TB of raw whole genome sequencing (WGS) data and 27 TB raw RNA-seq data into a series of geographically distributed data repositories, each running a uniform system for registering the data set, accepting and validating the raw read data and standardized metadata.

We utilized seven geographically distributed data repositories located at: (1) Barcelona Supercomputing Centre (BSC), (2) European Bioinformatics Institute (EMBL-EBI) in the UK, (3) German Cancer Research Center (DKFZ) in Germany; (4) the University of Tokyo in Japan; (5) Electronics and Telecommunications Research Institute (ETRI) in South Korea; (6) the Cancer Genome Hub (CGHub) and (7) the Bionimbus Protected Data Cloud (PDC) in the USA (Figure 2 and Suppl Table 1).

**Figure 2:**
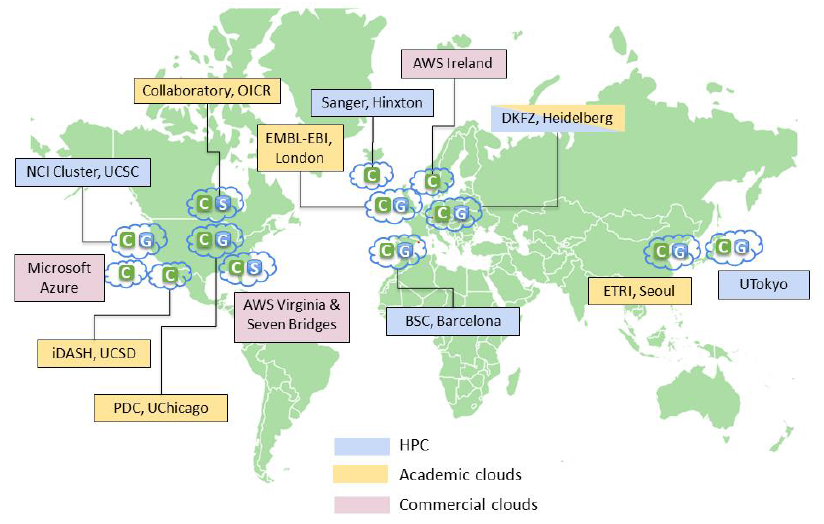
Geographical distribution of compute centers (C), GNOS servers (G), and S3-compatible data storage (S).

**Figure 3:**
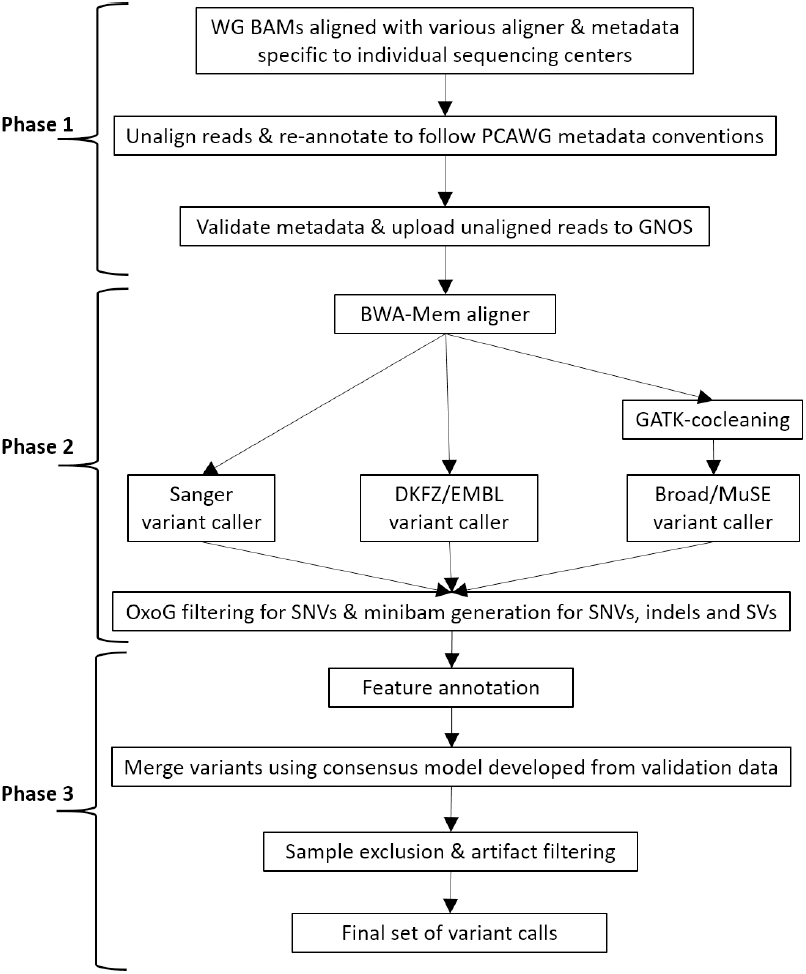
The uniform analysis of whole genomes involves three broad phases. Phase 1: Data marshalling and upload. Phase 2: Sequence alignment and variant calling. Phase 3: Variant merging and filtering. The algorithms for merging SNVs and indels are described in the PCAWG-1 paper, SVs in the PCAWG-6 paper, and CNVs in the PCAWG-11 paper.

To accept and validate sequence set uploads, each data repository ran a commercial software system, GNOS (Annai Systems). We chose GNOS because of the heavy testing it had previously received as the engine powering TCGA CGHub, and its support for validation of metadata according to the Sequence Read Archive (SRA) standard and file submission, strong user authentication and encryption, as well as its highly optimized data transfer protocol^6^. Each of the seven data centers initially allocated several hundred terabytes of storage to accept raw sequencing data from submitters within the region. The data centers also provided co-located compute resources to perform alignment and variant calling on the uploaded data.

Genomic data uploaded to the GNOS repositories was accompanied with detailed and accurate metadata to describe the cancer type, sample type, sequencing type and other attributes for managing and searching the files. We required that identifiers for project, donor, sample follow a standardized convention such that validation and auditing tools could be implemented. Most of the naming conventions in PCAWG were adopted from the well established ICGC data dictionary (http://docs.icgc.org/dictionary/about/).

Since most member projects at the time of upload already had sequencing reads aligned and annotated using their own metadata standards, a non-trivial effort was required to prepare the sequencing data for submission to GNOS. Each member project had to (1) prepare lane-level unaligned reads in BAM format, (2) reheader the BAM files with metadata following the PCAWG conventions, (3) generate metadata XML files, and (4) upload the BAM files along with the metadata XML files to GNOS. To facilitate this process, we developed the *PCAP-core* tool (https://github.com/ICGC-TCGA-PanCancer/PCAP-core) to extract the metadata from the BAM headers, validate the metadata, transform the metadata into the XML files conforming to the SRA specifications, and submitting the BAM files along with the metadata XML files to GNOS.

## Phase 2: Sequence Alignment and Variant Calling

We began the process of sequence alignment about two months after the uploading process had begun. Both tumor and matched normal reads were subjected to uniform sequence alignment using BWA-MEM^7^ on top of a common GRCh37-based reference genome that was enhanced with decoy sequences, viral sequences, and the revised Cambridge reference genome for the mitochondria.

Efforts by the project QC group demonstrated that employing multiple variant callers in ensemble fashion improved calling sensitivity^3^, thus the aligned tumor/normal pairs were subjected to somatic variant calling using three “best practice” software pipelines. These pipelines were developed by the Sanger Institute^8-11^; jointly by DKFZ^12^ and the European Molecular Biology Laboratory (EMBL)^13^; and the Broad Institute^14^ with contribution from MD Anderson Cancer Center-Baylor College of Medicine^15^. Each pipeline represents the best practices from the authoring organizations and include the current versions of each institute’s flagship tools. Each pipeline consists of multiple software tools for calling of single and multiple nucleotide variants (SNVs and MNVs), small insertions/deletions (indels), structural variants (SVs) and somatic copy number alterations (SCNAs). The minimum compute requirements, median runtime and the analytical algorithms for each pipeline are shown in Table 2.

**Table 2.** The five core workflows. Components for calling (1) SNVs, (2) indels, (3) SVs and (4) SCNAs in each of the three variant calling workflows are listed. Because we utilized a large number of compute environments with various configurations of cores and RAM, the average runtime for each pipelines varied with large standard deviations (Suppl Fig. 7-10). The runtime for the Broad pipeline included the 24 hours required to run GATK co-cleaning of BAMs. The measured runtime included time to download input files, but not the time to upload result files. (#) MuSE was developed at MD Anderson Cancer Center and Baylor College of Medicine.

|  | **BWA** | **Sanger** | **DKFZ/EMBL** | **Broad** | **OxoG** |
| --- | --- | --- | --- | --- | --- |
| Analytical components in workflow | BWA-Mem<br>Picard<br>Biobambam<br>samtools | CaVEMan<sup>1</sup><br>cgpPindel<sup>2</sup><br>BRASS<sup>3</sup><br>ascatNgs<sup>4</sup> | dkfz_snv<sup>1</sup><br>Platypus<sup>2</sup><br>DELLY<sup>3</sup><br>ACE-seq<sup>4</sup> | GATK cocleaning<br>MuTect<sup>1</sup><br>MuSE<sup>1,#</sup><br>Snowman<sup>2,3</sup><br>dRanger<sup>3</sup> | OxoG<br>VariantBam |
| Workflow controller | SeqWare | SeqWare | Roddy,<br>SeqWare | Galaxy | SeqWare |
| Recommended compute requirements | 4 cores,<br>15GB RAM | 16 cores,<br>4.5GB<br>RAM/core | 16 cores,<br>64GB RAM | 32 cores,<br>244GB RAM | 8 cores,<br>64GB RAM |
| Average runtime across all compute environments | 2.0 +/- 1.7<br>days | 5.3 +/- 5.5<br>days | 3.2 +/- 1.7<br>days | 5.1 +/- 2.2<br>days | 2.6 +/- 1.3<br>hours |
| Benchmark on AWS | 5.8 days on<br>4-core<br>m1.xlarge | 2.2 days on<br>32-core<br>r3.8xlarge | 1.7 days on<br>32-core<br>r3.8xlarge | 3.7 days on<br>32-core r3.8xlarge | 4 hours on<br>8-core<br>m2.4xlarge |
| Core hours per run | 557 | 1690 | 1306 | 2842 | 32 |
| Output files per run | 120GB | 2 GB | 5 GB | 35 GB | 1.5 GB |

When possible, both the alignment and variant calling pipelines were executed in the same regional compute centers to which the data sets were uploaded. As the project progressed, we utilized additional compute resources from AWS, Azure, iDASH, the Ontario Institute for Cancer Research (OICR), the Sanger Institute, and Seven Bridges (Figure 2). These centers computed on data sets located in the same region to optimize data transfer. Over the course of the project, some centers outpaced others and we rebalanced data sets as needed to use resources as efficiently as possible. Figure 1 shows the progress of the analytic pipelines with more details shown in Supplementary Figures 2-6.

## Phase 3: Variant merging, filtering, and synchronization

Following the completion of the three variant calling workflows, variants were passed to an additional pipeline referred as the “OxoG workflow”. This pipeline filtered out oxidative artifacts in SNVs using the OxoG algorithm^16^, normalized indels using the bcftools “norm” function, annotated genomic features for downstream merging of variants, and generated one “minibam” per specimen using the VariantBam algorithm^17^. Minibams are a novel format for representing the evidence that underlies genomic variant calls. Read pairs spanning a variant within a specified window were extracted from the whole genome BAM to generate the minibam. The windows we chose were +/- 10 base pairs (bp) for SNVs, +/- 200 bp for indels, and +/- 500 bp for SV breakpoints. The resulting minibams are about 0.5% of the size of whole genome BAMs, totalling to about four terabytes for all PCAWG specimens, making it much easier to download and store for the purpose of inspecting variants and their underlying read evidence.

Following filtering, we applied a series of merge algorithms to merge variants from the multiple variant calling pipelines into consensus call sets with higher accuracies than the individual pipelines alone. The SNV and indel merge algorithms were developed on the basis of experimental validation of the individual variant calling pipelines using deep targeted sequencing, a process detailed in the PCAWG-1 marker paper^4^. The algorithm for consensus SVs is described in the PCAWG-6 marker paper^18^. The consensus SCNAs were built upon the base-pair breakpoint results from the consensus SVs using a multi-tiered bespoke approach combining results from 6 SCNA algorithms^19^.

Following merging, the SNV, indel, SV and SCNA consensus call sets were subjected to intensive examination by multiple groups in order to identify anomalies and artefacts, including uneven coverage of the genome, strand and orientation bias, contamination with reads from non-human species, contamination of the library with DNA from an unrelated donor, and high rates of common germline polymorphisms among the somatic variant calls^4,11^. In keeping with our mission to provide a high-quality and uniformly annotated data set, we developed a series of filters to annotate and/or remove these artefacts. Tumor variant call sets that were deemed too problematic to use for downstream analysis were placed on an “exclusion list” (353 specimens, 176 donors). In addition, we established a “grey list” (150 specimens, 75 donors), of call sets that had failed some tests but not others and could be used, with caution, for certain types of downstream analysis. The criteria for classifying callsets into exclusion and grey list are described in more detail in the PCAWG-1 paper^10^.

Following the filtering steps, we used GNOS to synchronize the aligned reads and variant call sets among a small number of download sites for use by PCAWG downstream analysis working groups (Suppl Table 2). We also provided login credentials to members of PCAWG working groups for compute cloud-based access to the aligned read data across several of the regional data analysis centers, which avoided the overhead of downloading the data.

## Software and Protocols

This section describes the software and protocols developed for this project in more detail. All the software that we created for this project is available for use by any research group to conduct similar cloud-based cancer genome analyses economically and at scale.

### Centralized Metadata Management System

The metadata describing the donors, specimens, raw sequencing reads, WGS and RNA-Seq alignments, variant calls from the three pipelines, OxoG-filtered variants, and mini-BAMs were collected from globally distributed GNOS repositories, consolidated and indexed nightly using ElasticSearch (https://www.elastic.co) in a specially designed object graph model. This centrally managed metadata index was a key component of our operations and data provenance tracking. First, the metadata index was critical for tracking the status of each sequencing read set and for scheduling the next analytic step. The index also tracked the current location of each BAM and variant call set, allowing the pipelines to access the needed input data efficiently. Second, the metadata index provided the basis for a dashboard (http://pancancer.info) for all stakeholders to track day-to-day progress of each pipeline at each compute site. By reviewing the throughput of each compute site on a daily basis, we were able to identify issues early and to assign work accordingly to keep our compute resources productive. Third, the metadata index was also used by the ICGC Data Coordination Centre (DCC) to transfer PCAWG core datasets to long-term genomic data archive systems. Finally, the metadata index was imported into the ICGC Data Portal (https://dcc.icgc.org) to create a faceted search for PCAWG data allowing users to quickly locate data based on queries about the donor, cancer type, data type or data repositories.

### Docker Containers & Consonance

Given that the compute resources donated to the PCAWG project were a mix of cloud and HPC environments, we required a mechanism to encapsulate the analytical workflows to allow them to run smoothly across a wide variety of compute sites. The approaches we used evolved over time to incorporate better ways of abstracting and packaging tools to facilitate this portability. Initially, we used SeqWare workflow execution engine^20^ for bundling software and executing workflows, but this system required extensive and time consuming setup for the worker virtual machines (VMs). Later, we adopted Docker (http://www.docker.com) as a key enabling technology for running workflows in an infrastructure-independent manner. As a lightweight, infrastructure-agnostic containerization technology, Docker allowed PCAWG pipeline authors to fully encapsulate tools and system dependencies into a portable image. This included the fleet of VMs on commercial and academic clouds, as well as the project’s HPC clusters that were modified to support Docker containers. Each of our major pipelines was encapsulated in a single Docker image, along with a suitable workflow execution engine, reference data sets, and software libraries (Table 2).

Another key component of the PCAWG software infrastructure stack was cloud-agnostic technology to provision virtual machines on both academic and commercial clouds. Our initial attempts to scale the analytic pipelines across multiple cloud systems were complicated by transient failures in many of the academic cloud environments, subtle differences between seemingly identical clouds, and misconfigured services within the clouds. Initially, we attempted to replicate within the clouds standard components of conventional HPC environments, including shared file systems and cluster load balancing systems. However, we quickly learned that these perform poorly in the dynamic environments of the cloud. After several design iterations, we developed Consonance (https://github.com/consonance), a cloud-agnostic provisioning and queueing platform. For each of the cloud platforms in use in PCAWG, including OpenStack, VMWare, AWS, and Azure, Consonance provided a queue where work scheduling was decoupled from the worker nodes. As the fleet of working nodes shrank or expanded, each queue queried the centralized metadata index to obtain the next batch of tasks to execute. Consonance then created and maintained a fleet of worker VMs, launched new pipeline jobs, detected and relaunched failed VMs, and reran workflows as needed. Consonance allowed us to dynamically allocate cloud resources depending on the workload at hand, and even interacted with the AWS spot marketplace to minimize our commercial cloud costs.

### The Operations: whitelist, work queue, cloud shepherds

For the duration of the project, several personnel were required to operate the Docker images, Consonance and the metadata index effectively (Figure 4). Each compute environment was managed by a “cloud shepherd” responsible for completing the workflows on a set of pre-assigned donors or specimens. All the HPC environments (BSC, DKFZ, UTokyo, UCSC, Sanger) were shepherded by personnel local to the institute who were already familiar with the specific file systems and work schedulers, and obtained technical support from their local system administrators. The majority of the cloud environments (AWS, Azure, DKFZ, EMBL-EBI, ETRI, OICR, PDC) granted tenancy to OICR whose personnel acted as cloud shepherds. The other clouds (iDASH, SB), newly launched at the time, assigned their own cloud shepherds who also tested and fine tuned their environments in the process.

**Figure 4:**
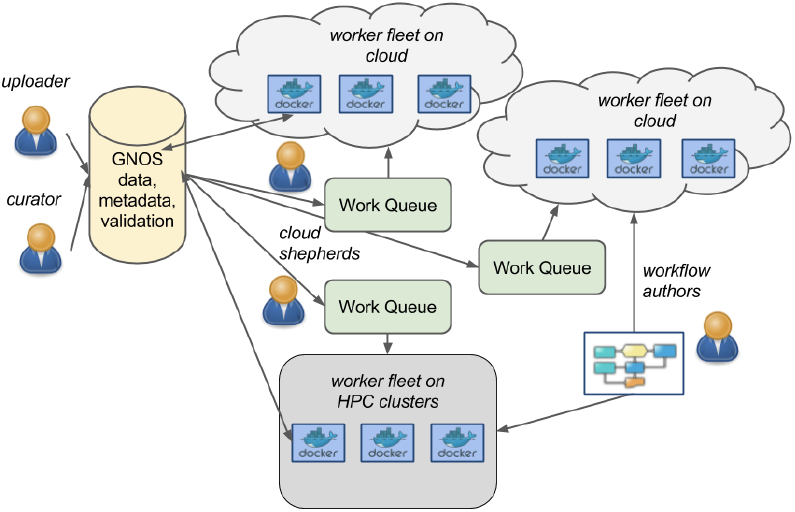
Infrastructure used on cloud and HPC compute environments for core analysis.

A project manager acted as the point of contact for all the cloud shepherds to report any technical issues and progress, such that the overall availability of compute resources and throughput at any time point could be estimated. Combining this knowledge with the information from the centralized metadata index, the project manager assigned donors and workflows to compute environments in the form of “whitelists” on a weekly basis. Cloud shepherds then added the whitelist of donors to their workflow queue for execution. This approach allowed us to be agile in responding to data availability disruptions, planned or unplanned downtime while optimizing data transfer and operations throughput.

While quotas shifted throughout the duration of the analysis, as demands and workloads on the individual centers changed, the overall peak commitment received was on the order of the 15,000 cores, approximately 60TB of RAM, and a peak usage of ~630 virtual machines.

### Software Distribution through Dockstore

The workflows used during PCAWG production include several PCAWG-specific elements that may limit their usability by researchers outside of the project. To facilitate the long term usage of these workflows by a broad range of cancer genomic researchers, we have simplified the tools to make most workflows standalone (Suppl Table 4). These Docker-packaged workflows have been extensively tested for their reproducibility and are registered on the Dockstore^21^ (http://dockstore.org), a service compliant with Global Alliance for Genomics and Health (GA4GH) standards to provide computational tools and workflows through Docker and described with Common Workflow Language^22^ (CWL). This enables other researchers to run the workflows on their own data, extend their utility, and replicate the work we have done in any CWL-compliant environment. By running the identical PCAWG workflows on their own data, researchers will be able to make direct comparisons and add to the existing PCAWG dataset.

The Docker-packaged BAM alignment and variant calling workflows were tested in different cloud environments and found to be easy to enact by third parties. Some discrepancies with the official data were observed and attributed to improvements in the underlying software (Sanger, Delly) or to the stochastic nature of the software, and deemed to have a low overall impact. Despite not achieving a completely identical results, the reproducibility of the process is satisfactory, especially considering that it involves software developed independently by different teams.

### Data Distribution / Data Portal

While GNOS was used for the core pipelines, Synapse^23^ was used to provide an interface to the files generated by the working groups and other intermediate results created throughout the project. Unlike GNOS which is focused on archival storage, Synapse allowed for collective editing in the form of a wiki, provenance tracking and versioning of results through a web interface as well as programmatic APIs. While Synapse provided an interface that allowed analyses to be shared rapidly across the consortia, the controlled access data was stored on a secure SFTP server provided by the National Cancer Institute (NCI). When the working groups complete their analysis, the metadata is retained in Synapse while the final version of the results is transferred to the ICGC Data Portal for archival.

In addition to GNOS-based repositories, the PCAWG dataset has been mirrored to multiple locations: the European Genome-phenome Archive (EGA, https://www.ebi.ac.uk/ega/studies/EGAS00001001692), AWS Simple Storage Service (S3, https://dcc.icgc.org/icgc-in-the-cloud/aws), and the Cancer Genome Collaboratory (http://cancercollaboratory.org). The data holdings at each repository at the time of publication are summarized in Suppl Table 2. To help researchers locate the PCAWG data, the ICGC Data Portal (https://dcc.icgc.org) provides a faceted search interface to query about donor, cancer type, data type or data repositories. Users can browse the collection of released PCAWG data and generate a manifest that facilitates downloading of the selected files.

The data repositories hosted at AWS S3 and the Collaboratory are powered by an open source object-based ICGC Storage System (https://github.com/icgc-dcc/dcc-storage) that enables fast, secure and multi-part downloads of files. Since AWS and the Collaboratory also have compute power co-located with the PCAWG data, they serve as effective cloud resources for researchers wishing to conduct further analyses on the PCAWG data without having to provision local compute resources and to download terabytes of data to their local compute environment.

## Discussion: Replicating PCAWG Analysis on Your Own Data

This project provided us with a rare opportunity to directly compare three categories of compute environment: traditional HPC, academic compute clouds and commercial clouds. In terms of stability and first time setup effort, we found that the traditional HPC environment routinely outperformed academic cloud systems, and often outperformed the commercial clouds. However, most of the academic cloud systems we worked with had been recently installed and some of the stability issues resulted from the shake-down period. The major benefit of the commercial clouds was the ability to scale compute resources up or down as needed, the ease of replicating the setup in different regions, and the availability of cloud-based data centers in different geographic regions, which allowed us to minimize data transfer overhead. For groups interested in replicating PCAWG results, or using the analytic pipelines for their own data, we are comfortable recommending running the analysis on a commercial cloud.

In terms of cost, we have summarized in Figure 5 the costs of computing on AWS and the tradeoff in accuracy if running a subset of the variant calling pipelines. The cost of aligning one normal specimen and one tumor specimen, and running three variant calling workflows followed by the OxoG workflow is about $100 per donor. This is based on a mean WGS coverage of 30X for normal specimens, and a bimodal coverage distribution with maxima at 38X and 60X for tumor specimens^24^. In addition, the hourly rate of the VMs are approximated from the spot instance pricing we experienced during production runs. With three variant calling workflows, we achieved an F1 score of 0.92. If one is willing to sacrifice some accuracy in order to reduce costs, then running only one variant calling workflow may be an option. Despite the higher costs, running two workflows does not result in increased accuracy. Unfortunately, we were not able to directly compare the analysis costs among commercial clouds, academic clouds and HPC due to the difficulty in assessing the fully loaded cost of provisioning and running an academic compute cluster.

**Figure 5:**
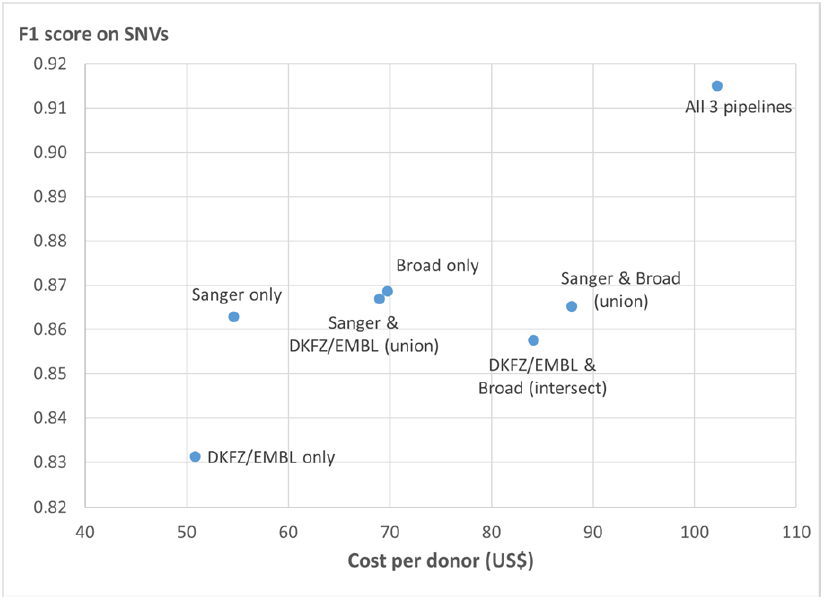
Costs for analyzing a tumor/normal pair through BWA-Mem, different combinations of variant calling pipelines, and OxoG filtering. The cost is calculated based on AWS instances at average spot pricing we experienced during the project, and includes egress costs to transfer the result files. PCAWG ran all 3 variant calling pipelines and achieved an F1 score of 0.9151 for SNVs. If running only one or two pipelines, there will be savings in cost but sacrifice in accuracy. Detailed cost analysis is shown in Suppl Table 3.

In terms of time, the major benefit of operating on commercial clouds is the availability of ample resources for simultaneous parallel runs. For example, in a scenario to analyze a total of 100 donors, one runs 200 VMs each aligning one tumor or normal specimen, followed by 300 VMs each running one of the three variant calling workflows on one donor, and 100 VMs to run OxoG workflow, the analysis will in principle take under 9 days to complete. In practice, additional time must be allowed for testing, scaling up, and the inevitability of failed jobs. A more realistic estimate of the time taken to run 100 donors through the complete PCAWG analysis on a commercial cloud is a few weeks.

Another issue when planning a large-scale genome analysis project is the variance in execution time from donor to donor. The variant calling pipelines took between 40 and 65 hours of wall time to complete a tumor/genome pair, with the EMBL/DKFZ pipeline running the quickest and the Broad and Sanger pipelines taking somewhat longer. In addition to the variant calling step, the Broad pipeline was preceded by a GATK co-cleaning process taking an additional 24 hours. For each pipeline there was significant variation in the runtime taken for each genome, and some tumor/normal pairs required an excessive amount of time to complete. Because long-running jobs can have economic and logistic impacts, we investigated the cause of this variation by applying linear regression to a number of features describing the raw sequencing sets, including coverage, read quality and mapping scores, number of mismatched end pairs and others (data not shown). We found that a single factor, genomic coverage, explained the variation in wall clock time which increased roughly linearly with coverage.

In conclusion, we tackled the challenge of performing uniform analysis on a large dataset across a geographically and technologically disparate collection of compute resources by developing technologies that realized the efficiencies of moving algorithms to the data. This is becoming a necessity as genomic datasets continue to increase in size and are geographically distributed with some jurisdictions restricting the geographical storage and computing of specific datasets. Our approach serves as a model for large scale collaborative efforts that engage many organizations and spread the computation work around the globe.

Our effort resulted in three key deliverables. First and foremost, we produced a high-quality, validated consensus variant and alignment dataset of 2,834 cancer donors. To date, this is the largest whole genome cancer dataset analyzed in a consistent and uniform way. The dataset formed the basis for the research by the PCAWG working groups, and will continue to provide value to the research community for many years into the future. Second, we produced a series of best-practice analytical workflows that are portable through the use of Docker and are available on the Dockstore. These workflows are usable in a multitude of compute environments giving researchers the ability to replicate our analysis on their own data. Finally, the infrastructure we built to coordinate analyses between cloud and HPC environments will be helpful for other projects requiring the same distributed approaches.

## Acknowledgements

The authors would like to acknowledge the donation of the following compute resources: the PRACE Research Infrastructure resource MareNostrum3 at Barcelona Supercomputing Center with technical expertise provided by the Red Española de Supercomputación and funding support by the Spanish Ministry of Health, ISCIII, in the project Instituto Nacional de Bioinformática (PRB2: PT13/0001/0028); the Cancer Genome Collaboratory, jointly funded by the Natural Sciences and Engineering Research Council of Canada, the Canadian Institutes of Health Research, Genome Canada, and the Canada Foundation for Innovation, and with in-kind support from the Ontario Research Fund of the Ministry of Research, Innovation and Science through the Discovery Frontiers: Advancing Big Data Science in Genomics Research program (grant no. RGPGR/448167-2013); the EMBL-EBI Embassy Cloud supported by UK’s (BBSRC) Large Facilities Capital Fund and Cancer Research UK’s EMBL-EBI Bioinformatics Resource (grant no. C32939/A20952); sFTP server provided by the Center for Biomedical Informatics & Information Technology (CBIIT) at National Cancer Institute; infrastructure at the Ontario Institute for Cancer Research funded by the Government of Ontario and the Canada Foundation for Innovation (Project #21039); ETRI’s OpenStack supported by Institute for Information & communications Technology Promotion with funding from the Korea government (MSIP) (No.B0101-15-0104, The Development of Supercomputing System for the Genome Analysis), Ministry of Health & Welfare, Republic of Korea (grant no: HI14C0072), Korean national research foundation (grant no NRF-2017R1A2B2012796, NRF-2016R1D1A1B03934110), and generous support from Wan Choi and Kwang-Sung; ‘Shirokane’ provided by Human Genome Center, the Institute of Medical Science, the University of Tokyo along with technical assistance from Hitachi, Ltd.; Microsoft Azure contributed through a grant to the UC Santa Cruz Genomics Institute and supported by the National Human Genome Research Institute of the National Institutes of Health (grant no U54HG007990) and NCI ITCR (grant no 1R01CA180778); iDASH HIPAA cloud which is a member of the NIH/NHLBI National Centers for Biomedical Computing (U54HL108460) to UC San Diego Health Sciences, Department of Biomedical Informatics.

In addition, the Broad team was supported by G.G. funds at MGH and Broad Institute. The DKFZ team was supported by the BMBF-funded Heidelberg Center for Human Bioinformatics (HD-HuB) within the German Network for Bioinformatics Infrastructure (de.NBI) (#031A537A, #031A537C) and the BMBF-funded grants ICGC PedBrain (01KU1201A, 01KU1201B), ICGC EOPC (01KU1001A), ICGC MMML-seq (01KU1002B), and ICGC DE-MINING (01KU1505E). Variant calling with the DKFZ/EMBL pipeline made use of the Roddy framework, and provision of data and metadata of the German ICGC projects was assisted by the One Touch Pipeline (OTP). The OICR team was funded by the Government of Ontario and the Canada Foundation for Innovation (Project #21039). The Sanger team was supported by the Wellcome Trust grant (098051) with contributions by Shriram G Bhosle, David R Jones, Andrew Menzies, Lucy Stebbings, Jon W Teague.

## Supplementary Information

**Supplementary Figure 1:**
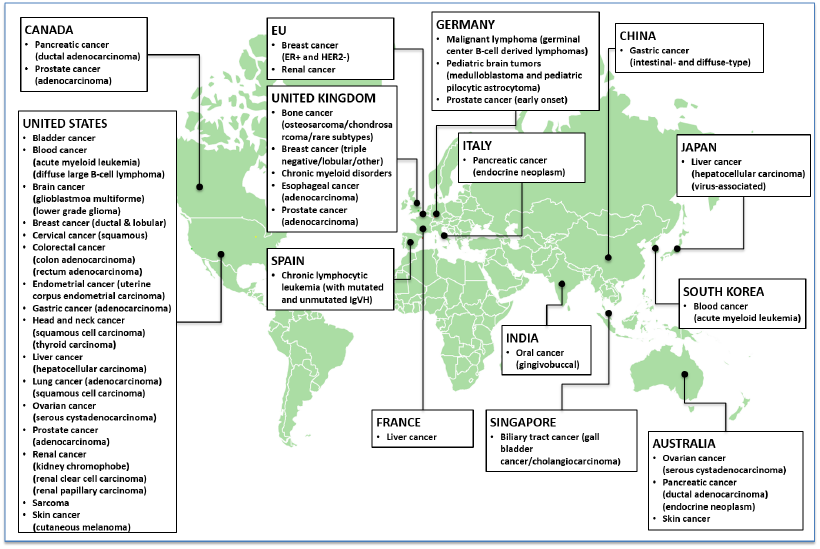
Whole genomes from 2,834 donors across 39 cancer types were collected from 48 ICGC and TCGA projects in 14 jurisdictions.

**Supplementary Figure 2:**
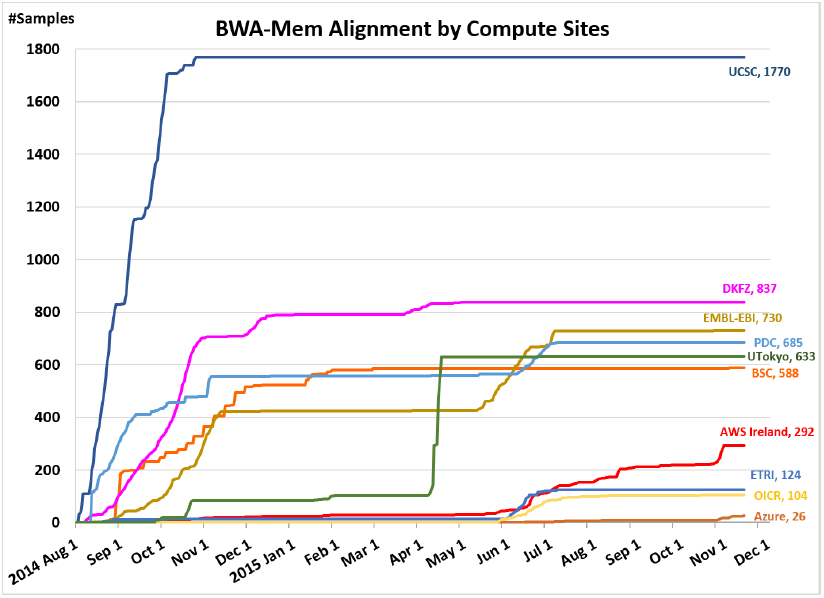
Progress of BWA-Mem alignment over time at 7 compute sites.

**Supplementary Figure 3:**
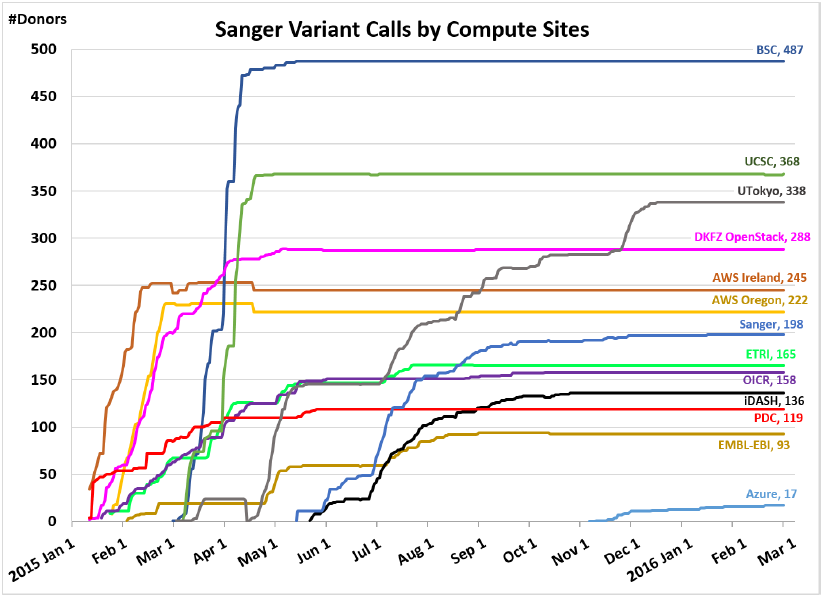
Progress of Sanger variant calling workflow over time at 13 compute sites.

**Supplementary Figure 4:**
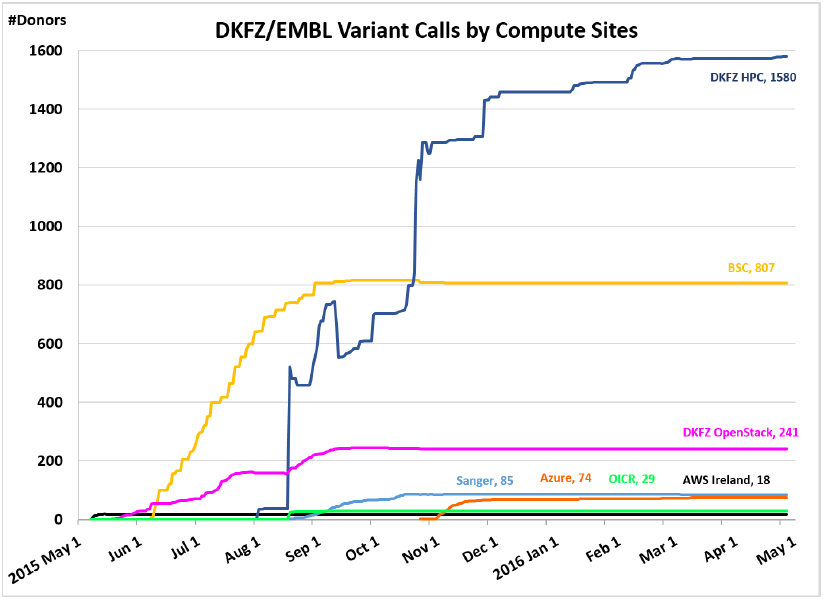
Progress of DKFZ/EMBL variant calling workflow over time at 7 compute sites.

**Supplementary Figure 5:**
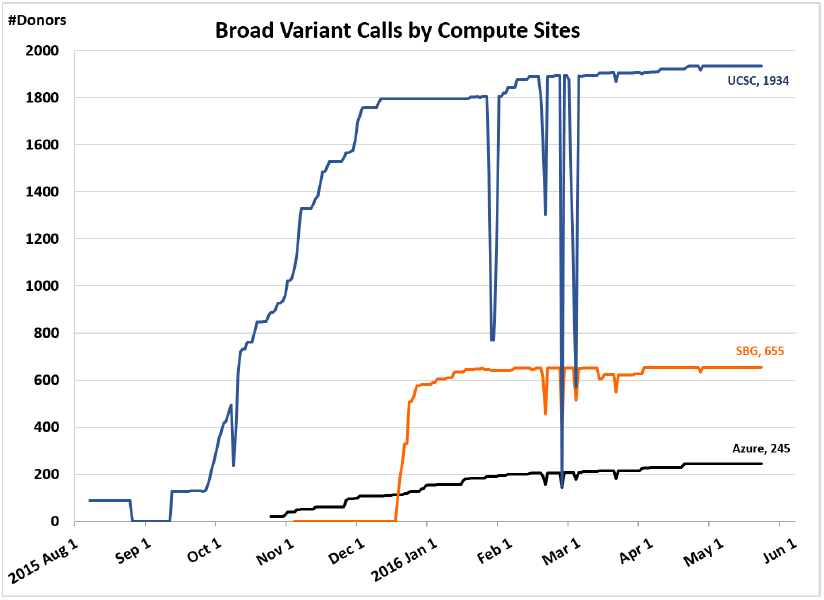
Progress of Broad variant calling workflow over time at 3 compute sites.

**Supplementary Figure 6:**
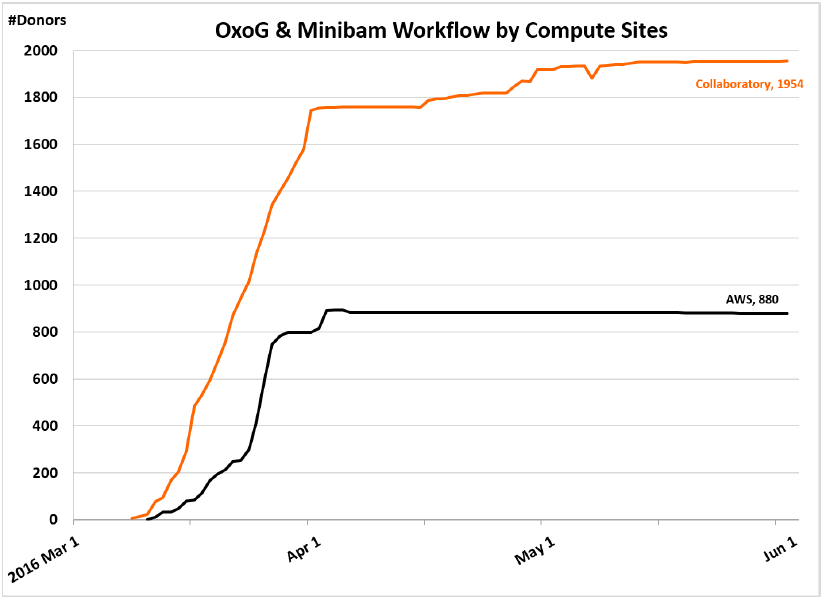
Progress of OxoG and minibam workflow over time at 2 compute sites.

**Supplementary Figure 7:**
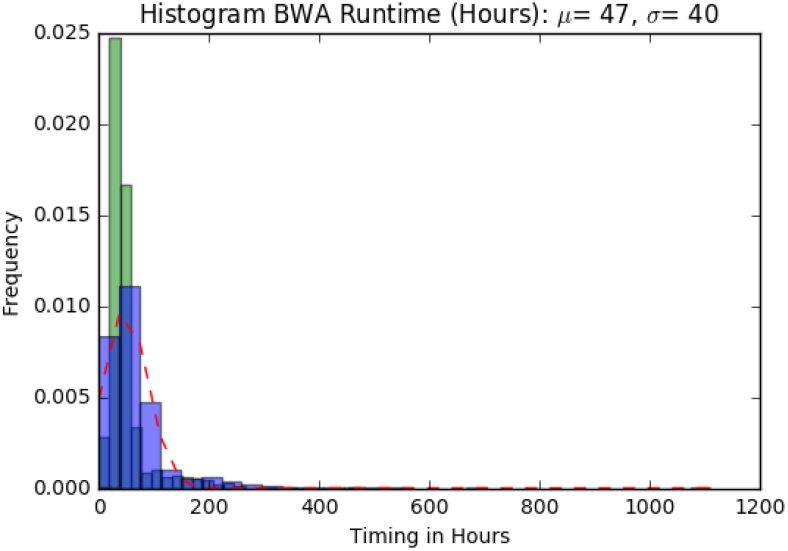
Average runtimes for BWA-Mem alignment workflow

**Supplementary Figure 8:**
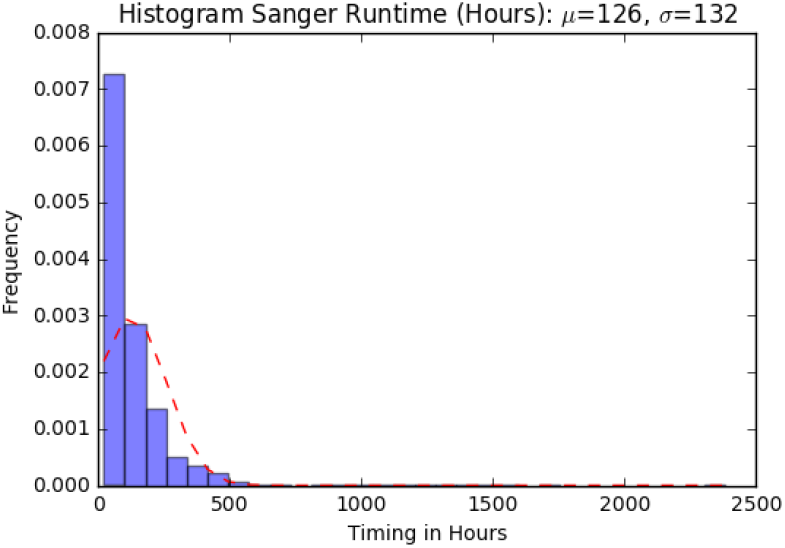
Average runtime for the Sanger somatic variant calling workflow.

**Supplementary Figure 9:**
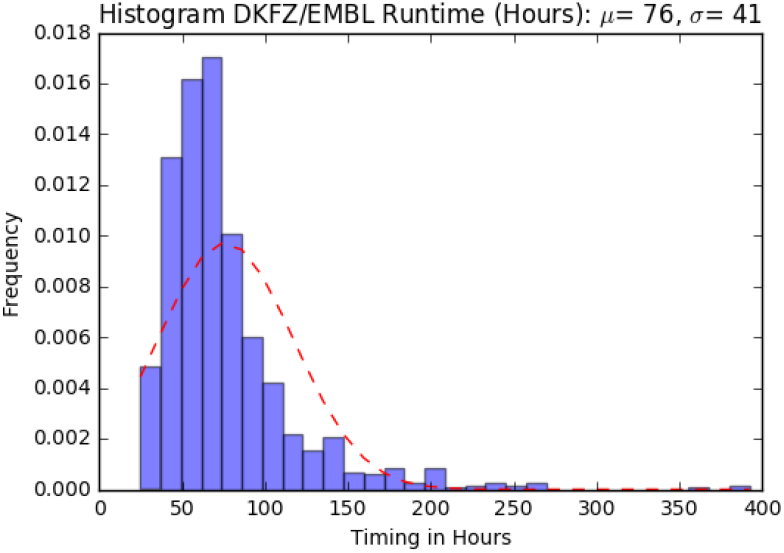
Average runtime for the DKFZ/EMBL somatic variant calling workflow.

**Supplementary Figure 10:**
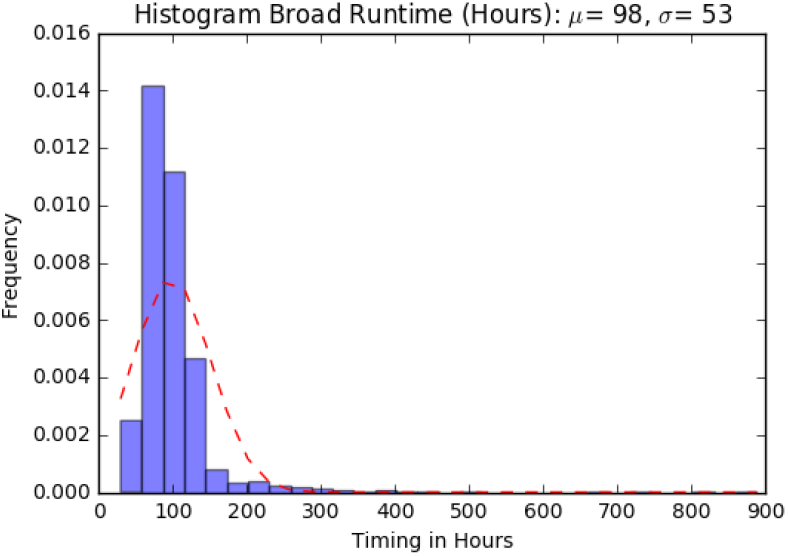
Average runtime for the Broad somatic variant calling workflow. Preceding the variant calling workflow, the GATK co-cleaning step takes an additional 24 hours.

**Supplementary Table 1.** Percentage samples/donors run at each site for each pipeline

|  | **BWA** | **Sanger** | **DKFZ/EMBL** | **Broad/MuSE** | **OxoG** |
| --- | --- | --- | --- | --- | --- |
| **AWS Ireland** | 5.0 | 16.4 | 0.6 |  | 31.1 |
| **Azure** | 0.4 | 0.6 | 2.6 | 8.6 |  |
| **BSC** | 10.2 | 17.2 | 28.5 |  |  |
| **Collaboratory** |  |  |  |  | 68.9 |
| **DKFZ (HPC)** |  |  | 55.8 |  |  |
| **DKFZ (OpenStack)** | 14.5 | 10.2 | 8.5 |  |  |
| **EMBL-EBI** | 12.6 | 3.3 |  |  |  |
| **ETRI** | 2.1 | 5.8 |  |  |  |
| **iDASH** |  | 4.8 |  |  |  |
| **OICR** | 1.8 | 5.6 | 1.0 |  |  |
| **PDC** | 11.8 | 4.2 |  |  |  |
| **Sanger** |  | 7.0 | 3.0 |  |  |
| **Seven Bridges** |  |  |  | 23.1 |  |
| **UCSC** | 30.6 | 13.0 |  | 68.2 |  |
| **UTokyo** | 10.9 | 11.9 |  |  |  |

**Supplementary Table 2.** Data distribution as of May 2017. While ETRI GNOS and CGHub served as data centres during the project, they have since been retired. Variant calls include those from individual variant calling pipelines and the final consensus callsets. Long-term repositories are denoted by asterisk (*) and will increase their data holdings over time while GNOS servers are gradually being retired. Latest information can be found at https://dcc.icgc.org/repositories

|  | ICGC Data |  |  | TCGA Data |  |  |
| --- | --- | --- | --- | --- | --- | --- |
| Data Repository | % WG Alignments (534 TB) | % RNA-Seq Alignments (13 TB) | % Variant calls (520 GB) | % WG Alignments (240 TB) | % RNA-Seq Alignments (14 TB) | % Variant calls (228 GB) |
| BSC GNOS | 100.0 | 30.0 | 0.3 |  |  |  |
| DKFZ GNOS | 25.0 |  | 62.9 |  |  |  |
| EMBL-EBI GNOS | 100.0 | 59.3 | 98.6 |  |  |  |
| UTokyo GNOS | 54.6 | 17.1 | 1.6 |  |  |  |
| UChicago-ICGC GNOS | 16.8 | 40.3 | 28.7 |  |  |  |
| UChicago-TCGA GNOS |  |  |  | 100.0 | 100.0 | 100.0 |
| EGA* | 97.8 |  |  |  |  |  |
| Collaboratory* | 100.0 | 100.0 | 100.0 |  |  |  |
| AWS* | 76.7 | 80.1 | 75.1 |  |  |  |
| Bionimbus PDC* |  |  |  | 100.0 | 100.0 | 0.2 |

The following set of tables show how costs are calculated for Figure 5 which compares the costs and accuracies of running the different combination of variant calling pipelines.

**Supplementary Table 3a.**
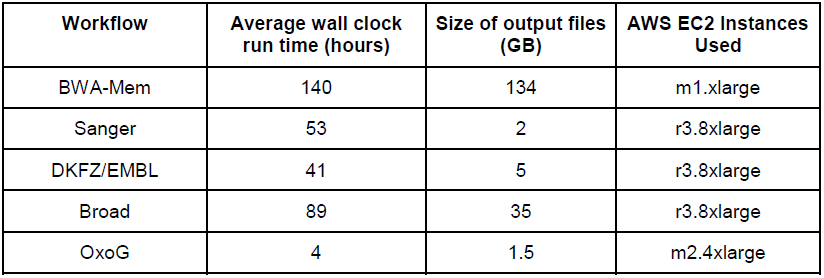
The average run time for each workflow was rounded up to the nearest hour to reflect how AWS charges for EC2 instances that run for part of an hour. The size of the output files are noted as they contribute to either egress or storage costs.

**Supplementary Table 3b.** The project utilized EC2 spot instances in US East (N. Virginia), US West (Oregon), EU (Ireland) regions. Because spot pricing fluctuates, users should consult real-time information. The average spot pricing listed here was based on our own usage throughout the project.

| AWS EC2 Instances | vCPU | Mem (GiB) | Storage (GB) | Average spot pricing |
| --- | --- | --- | --- | --- |
| m1.xlarge | 4 | 15 | 4 x 420 | \$0.0426 |
| r3.8xlarge | 32 | 244 | 2 x 320 | \$0.3382 |
| m2.4xlarge | 8 | 68.4 | 2 x 840 | \$0.0834 |

**Supplementary Table 3c.** Cost calculations are based on the above spot pricing and an egress cost of $0.09 per GB. The analysis time is made up of 3 steps: (1) running the BWA-Mem workflow on two separate instances to align simultaneously one tumor and one normal specimen; (2) running the variant calling workflows simultaneously with the longest running workflow dictating the run time of this step; (3) running the OxoG workflow after all variant calling workflows are completed. If analyzing 100 donors with all 3 variant calling pipelines, the analysis will involve running a fleet of 200, 300 and 100 EC2 instances, respectively in the 3 steps. We have no other significant storage cost as the reference files amount to ~35GB costing under $1/month in S3. An alternative to transferring the data out is to store the 312 GB of data for each donor in S3 for under $8/month.

| Variant Calling Pipelines | Total Cost | Compute Cost | Egress Cost | Analysis Time (days) | Median Sensitivity, Precision, F1 |
| --- | --- | --- | --- | --- | --- |
| All 3 pipelines | 102.19 | 7.15 | 28.04 | 9.7 | 0.9047 +/- 0.03145<br>0.9348 +/- 0.03785<br>0.9151 +/- 0.02820 |
| Sanger only | 54.63 | 30.19 | 24.44 | 8.2 | 0.8032 +/- 0.06515<br>0.9550 +/- 0.03855<br>0.8629 +/- 0.04795 |
| DKFZ/EMBL only | 50.84 | 26.13 | 24.71 | 7.7 | 0.7565 +/- 0.0544<br>0.9352 +/- 0.0365<br>0.8313 +/- 0.05125 |
| Broad only | 69.77 | 42.36 | 27.41 | 9.7 | 0.9095 +/- 0.01955<br>0.8386 +/- 0.06335<br>0.8687 +/- 0.04085 |
| Sanger & DKFZ/EMBL | 68.94 | 44.05 | 24.89 | 8.2 | <u>Union</u><br>0.8454 +/- 0.0572<br>0.9032 +/- 0.04405<br>0.8669 +/- 0.0509<br><u>Intersect</u><br>0.7228 +/- 0.05385<br>0.9954 +/- 0.00980<br>0.8216 +/- 0.04390 |
| Sanger & Broad | 87.88 | 60.29 | 27.59 | 9.7 | <u>Union</u><br>0.9374 +/- 0.01935<br>0.8183 +/- 0.06395<br>0.8653 +/- 0.04220<br><u>Intersect</u><br>0.7856 +/- 0.0566<br>0.9913 +/- 0.0111<br>0.8632 +/- 0.03755 |
| DKFZ/EMBL & Broad | 84.09 | 56.23 | 27.86 | 9.7 | <u>Union</u><br>0.9339 +/- 0.01955<br>0.801 +/- 0.06505<br>0.8576 +/- 0.0429<br><u>Intersect</u><br>0.7384 +/- 0.05865<br>0.9939 +/- 0.0186<br>0.8315 +/- 0.0456 |

**Supplementary Table 4.** DOIs for PCAWG core analysis workflows

| Workflow/Tool | Dockstore | Latest DOI | Version | Github |
| --- | --- | --- | --- | --- |
| pcawg-bwa-mem-workflow | https://dockstore.org/containers/quay.io/pancancer/pcawg-bwa-mem-workflow | https://doi.org/10.5281/zenodo.192377 | 2.6.8_1.2 | https://github.com/ICGC-TCGA-PanCancer/Seqware-BWA-Workflow |
| pcawg-dkfz-workflow | https://dockstore.org/containers/quay.io/pancancer/pcawg-dkfz-workflow | https://doi.org/10.5281/zenodo.192376 | 2.0.1_cwl1.0 | https://github.com/ICGC-TCGA-PanCancer/DEWrapperWorkflow |
| pcawg-sanger-cgp-workflow | https://dockstore.org/containers/quay.io/pancancer/pcawg-sanger-cgp-workflow | https://doi.org/10.5281/zenodo.192162 | 2.0.3 | https://github.com/ICGC-TCGA-PanCancer/CGP-Somatic-Docker |
| pcawg_delly_workflow | https://dockstore.org/containers/quay.io/pancancer/pcawg_delly_workflow | https://doi.org/10.5281/zenodo.192166 | 2.0.1-cwl1.0 | https://github.com/ICGC-TCGA-PanCancer/DEWrapperWorkflow |
| broad |  |  |  |  |
| oxog |  |  |  |  |

## References

1. Network, T.C.G.A.R. et al. The Cancer Genome Atlas Pan-Cancer analysis project. Nature Genetics 45, 1113–1120 (2013).

2. PCAWG-3. Pan-Cancer Study of Recurrent and Heterogeneous RNA Aberrations and Association with Whole-Genome Variants. (in preparation).

3. Alioto, T.S. et al. A comprehensive assessment of somatic mutation detection in cancer using whole-genome sequencing. Nat Commun 6, 10001 (2015).

4. PCAWG-1. Consistent Detection of Short Somatic Mutations in 2,778 Cancer Whole Genomes. (in preparation).

5. Phillips, M. & Knoppers, B. Building an International Code of Conduct for Genomic Cloud Research. (in preparation).

6. Wilks, C. et al. The Cancer Genomics Hub (CGHub): overcoming cancer through the power of torrential data. Database (Oxford) 2014(2014).

7. Li, H. Aligning sequence reads, clone sequences and assembly contigs with BWA-MEM. (2013).

8. Jones, D. et al. cgpCaVEManWrapper: Simple Execution of CaVEMan in Order to Detect Somatic Single Nucleotide Variants in NGS Data. Curr Protoc Bioinformatics 56, 15.10.1-15.10.18 (2016).

9. Raine, K.M. et al. cgpPindel: Identifying Somatically Acquired Insertion and Deletion Events from Paired End Sequencing. Curr Protoc Bioinformatics 52, 15.7.1-12 (2015).

10. Raine, K.M. et al. ascatNgs: Identifying Somatically Acquired Copy-Number Alterations from Whole-Genome Sequencing Data. Curr Protoc Bioinformatics 56, 15.9.1-15.9.17 (2016).

11. BRASS. (https://github.com/cancerit/BRASS).

12. Rimmer, A. et al. Integrating mapping-, assembly- and haplotype-based approaches for calling variants in clinical sequencing applications. Nat Genet 46, 912–8 (2014).

13. Rausch, T. et al. DELLY: structural variant discovery by integrated paired-end and split-read analysis. Bioinformatics 28, i333–i339 (2012).

14. Cibulskis, K. et al. Sensitive detection of somatic point mutations in impure and heterogeneous cancer samples. Nat Biotechnol 31, 213–9 (2013).

15. Fan, Y. et al. MuSE: accounting for tumor heterogeneity using a sample-specific error model improves sensitivity and specificity in mutation calling from sequencing data. Genome Biol 17, 178 (2016).

16. Costello, M. et al. Discovery and characterization of artifactual mutations in deep coverage targeted capture sequencing data due to oxidative DNA damage during sample preparation. Nucleic Acids Res 41, e67 (2013).

17. Wala, J., Zhang, C.Z., Meyerson, M. & Beroukhim, R. VariantBam: filtering and profiling of next-generational sequencing data using region-specific rules. Bioinformatics 32, 2029-31 (2016).

18. PCAWG-6. PCAWG-6 paper. (in preparation).

19. PCAWG-11. PCAWG-11 paper. (in preparation).

20. O’Connor, B.D., Merriman, B. & Nelson, S.F. SeqWare Query Engine: storing and searching sequence data in the cloud. BMC Bioinformatics 11 Suppl 12, S2 (2010).

21. O’Connor, B.D. et al. The Dockstore: enabling modular, community-focused sharing of Docker-based genomics tools and workflows. F1000Res 6, 52 (2017).

22. Amstutz, P. et al. Common Workflow Language, v1.0. figshare (2016).

23. Omberg, L. et al. Enabling transparent and collaborative computational analysis of 12 tumor types within The Cancer Genome Atlas. Nat Genet 45, 1121–6 (2013).

24. PCAWG-QC. Framework for quality assessment of whole genome, cancer sequences. (in preparation).

